# Insomnia score: predictive ability of insomnia, high-diagnostic and prognostic value for cancer

**DOI:** 10.1101/2024.08.13.607749

**Authors:** Dan Xiao, Yixin Lan, Haohan Zhang, Siyu Li, Chuandong Zhou

**Author notes:** Address correspondence to Hongwei Shao,. The e-mail address of each author(in the order of authorship),. The author order was determined by the corresponding author after discussion.

## Abstract

**Objective:** This study utilized bioinformatics methods to investigate the association between insomnia and cancer.

**Methods:** We identified key insomnia-related genes and constructed an insomnia score (riskScore = ∑ Expression_gene_ * Coefficient) based on an insomnia dataset using differential analysis, Weighted Gene Co-expression Network Analysis (WGCNA), and Least Absolute Shrinkage and Selection Operator (LASSO) regression. The predictive ability of the insomnia score was validated in the insomnia dataset using Receiver Operating Characteristic (ROC) analysis. Finally, the diagnostic and prognostic value of the insomnia score was assessed in a pan-cancer cohort.

**Results:** Differential analysis, WGCNA, and LASSO screening of the insomnia dataset yielded three key insomnia-related genes (PTMA, NLRP8, and CCBE1). An insomnia score was then constructed using the formula: riskScore = -0.8039667 * Expression_PTMA_ + - 3.1975230 * Expression_NLRP8_ + -0.4957560 * Expression_CCBE1_. The ROC analysis demonstrated that the insomnia score has excellent predictive ability for insomnia. Furthermore, pan-cancer analysis illustrated the diagnostic and prognostic value of the insomnia score across various cancers.

**Conclusion:** We successfully constructed an insomnia score and demonstrated its diagnostic and prognostic value in multiple cancers.

## 1. Introduction

In modern daily life, people are often plagued by insomnia[1], and persistent or chronic insomnia is considered a brain disorder[2]. In clinical practice, insomnia is defined as a disorder characterized by difficulty falling asleep or maintaining sleep with daytime symptoms, with a prevalence of 10% to 20%[3]. The older age group is predominant among those who suffer from insomnia[4], but insomnia is also prevalent in the adolescent population[5]. Insomnia is also often associated with an unhealthy state of the body[6], and a number of studies have pointed to a correlation or causal relationship between insomnia and diseases such as depression[7,8], hypertension[9,10], coronary heart disease[11], metabolic syndrome[12], and diabetes mellitus[13]. Indeed, two phenotypes of insomnia exist, one associated with physiologic hyperarousal and concomitant disorders or sequelae of other diseases, and the other with cognitive-emotional and cortical arousal[14].

In cancer patients, insomnia is usually a complication of cancer[15–20], and the process of treating cancer often leads to insomnia in cancer patients [17,21–23]. Several recent studies have shown that insomnia leads to an increased risk of cancer[24–29], therefore, it is of great clinical importance to study the association between insomnia and cancer, but there are no studies exploring the association between insomnia and cancer based on gene microarray data.

Based on gene expression data, many studies have used bioinformatics methods including WGCNA[30] and LASSO[31] to select signature genes[32–36] or construct scoring models[35,37], and the results have shown good reliability and accuracy. This study synthesized previous studies using methods including WGCNA and LASSO to both select signature genes and construct model scores.

This study was based on published insomnia microarray data (GSE208668 [38] and GSE40562 [39]) on the GEO (https://www.ncbi.nlm.nih.gov/) database website and UCSC Xena (https://xenabrowser.net/datapages/) [40] provide pan-cancer data to assess the diagnostic and prognostic value of insomnia in cancer. Briefly, first, we identified insomnia candidate genes in the GSE208668 dataset using differential analysis and WGCNA, then screened insomnia core genes in combination with the LASSO algorithm and constructed insomnia score using the coefficients provided by the LASSO algorithm, and then we validated the ability of insomnia score in GSE208668 and GSE40562 to predict insomnia. Finally, we calculated the insomnia score in the TCGA pan-cancer cohort and the GDC Pan-Cancer cohort at UCSC Xena and evaluated its diagnostic and prognostic value in cancer. The results showed that insomnia score had excellent ability to predict insomnia in GSE208668 and GSE40562, and insomnia score had high diagnostic and prognostic value in a variety of cancers, which implies that insomnia has a strong association with the development of cancer and prognosis. In conclusion, this study developed an insomnia score with good predictive ability for insomnia, and the broad diagnostic and prognostic value of this score in cancer was determined by pan-cancer analysis. This work emphasizes the diagnostic and prognostic value of insomnia in a wide range of cancers, providing an important reference and a new perspective for research on the association between insomnia and cancer.

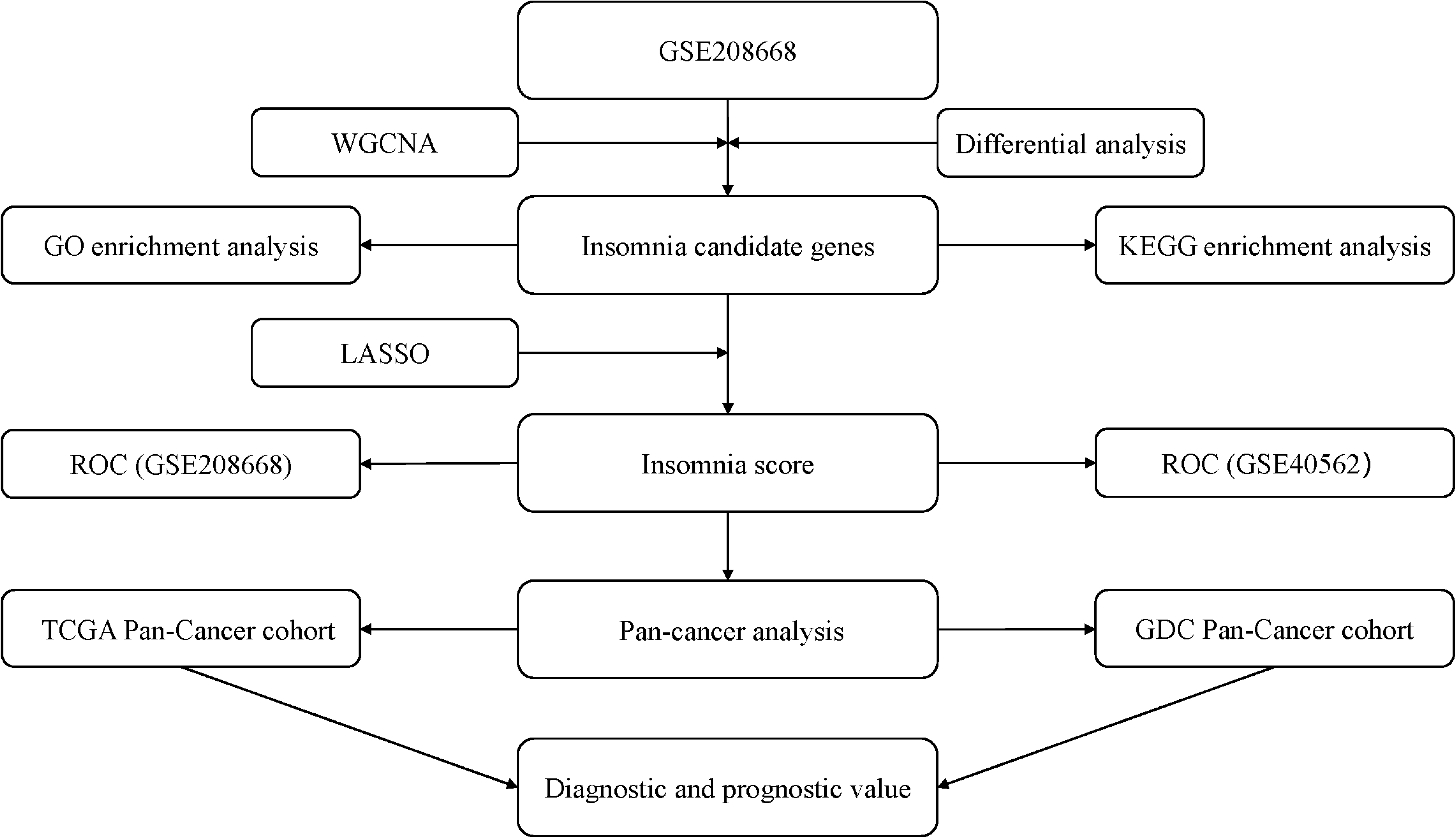
Graphical summary of this study. The arrows point to represent the workflow of the study.

## 2. Materials and methods

### 2.1 Data acquisition and processing

Insomnia-related datasets GSE208668 and GSE40562 were downloaded from the GEO (https://www.ncbi.nlm.nih.gov/) database using the R package GEOquery[41], and then the probe microarray data were converted to gene microarray data using the platform annotation file.GSE208668 was a collection of gene expression data from 17 individuals with insomnia in GSE208668 is a collection of gene expression data of peripheral blood mononuclear cells from 17 elderly people with insomnia and 25 elderly people without insomnia.GSE40562 is a collection of microarray data of thalamus or parietal lobe from 6 patients with fatal familial insomnia and normal people. Genes with no expression values in some samples were removed.

We downloaded TPM-type gene expression data for the TCGA Pan-Cancer cohort from UCSC Xena (https://xenabrowser.net/datapages/), which includes 10,535 samples, for conducting the TCGA pan-cancer study. Additionally, we downloaded curated clinical data for the TCGA Pan- Cancer cohort, which includes overall survival data, for the pan-cancer analysis.

The GDC Pan-Cancer cohort at UCSC Xena contains gene expression data from both the TCGA program and the TARGET program. Upon inspection, the samples from the TCGA program in the GDC Pan-Cancer cohort were different from those in the TCGA Pan-Cancer cohort (Supplementary file1). The gene expression data of the FPKM type from the GDC Pan-Cancer cohort, comprising 11,768 samples, was downloaded for conducting the Pan-Cancer study. Survival data from the GDC Pan-Cancer cohort, including overall survival data, were also downloaded as clinical data for the Pan-Cancer analysis.

### 2.2 Differential analysis

Gene symbols shared among GSE208668, GSE40562, and the TCGA Pan- Cancer cohort were screened from GSE208668 for comparison between insomnia and non-insomnia groups. Differential expression analysis was conducted using the limma software package. The criteria for identifying significantly different genes were set as log2FoldChange > 2 and FDR < 0.05.

### 2.3 WGCNA

After screening the gene symbols shared by GSE208668, GSE40562, and TCGA Pan-Cancer in GSE208668, we further selected gene symbols with higher variance in the gene expression matrix. Specifically, we retained gene symbols with a median absolute deviation higher than the first quartile and the 0.01 maximum for subsequent WGCNA analysis. The WGCNA analysis was performed using the WGCNA package[30], generating a weighted adjacency matrix with a soft threshold of 27 and evaluating the scale-free topology with a scale-free R^2^ = 0.85. Modules were hierarchically clustered based on the computed feature genes, each containing at least 50 genes (minModuleSize = 50), and similar modules were merged with a 75% similarity threshold (mergeCutHeight = 0.25). In this study, we calculated correlations between modules and clinical data, identified MEblue as a key insomnia-associated module, and conducted subsequent analyses with genes within the MEblue module.

### 2.4 GO and KEGG enrichment analysis

The significantly different genes identified in GSE208668 and the genes within the MEblue module were intersected. The intersected genes were then subjected to GO and KEGG enrichment analysis using the clusterProfiler[42] and enrichplot packages. GO terms or KEGG pathways that were significantly enriched were identified with a threshold of FDR < 0.05. The enrichment results were visualized using cnetplot. When more than five significantly enriched results were identified, the result with the smallest FDR was displayed; if the FDR was identical, the result with the highest number of genes covered by terms or pathways was shown. If fewer than five significantly enriched results were identified, all significantly enriched results were displayed.

### 2.5 Construction and validation of an insomnia score

Intersecting gene expressions of significantly different genes and genes within the MEblue module were obtained at GSE208668, and LASSO analysis was performed using the glmnet package[31] to screen for insomnia key genes. Insomnia score were obtained according to the riskScore formula: riskScore=∑Expression_gene_*Coefficient(expression is the average expression of a gene and coefficient are generated by LASSO regression). ROC analyses were performed using the pROC package[43] to determine the reliability of insomnia score and insomnia core genes in predicting insomnia symptoms in GSE208668 and dataset GSE40562, with an AUC >0.7 considered to have excellent predictive performance.

### 2.6 Pan-cancer analysis of the TCGA Pan-Cancer cohort

Gene expression data from samples with shared clinical data were screened in the TCGA Pan-Cancer cohort. The insomnia score for the shared samples was calculated according to the risk score formula, and differences in insomnia scores between cancer samples and normal samples in different cancers were analyzed using the Wilcoxon rank-sum test (P-value < 0.05 was considered statistically significant). Cancers with significant differences in insomnia scores between normal and cancer tissues were selected, and the performance of the insomnia score in diagnosing cancer was assessed by ROC analysis using the pROC package, with AUC > 0.7 considered indicative of high performance and diagnostic value.

Univariate Cox proportional hazards regression analysis was performed using the survival package to determine the correlation between insomnia and overall survival in tissue samples of 33 cancers. The results were presented in a forest plot. For those cancers with a significant correlation between insomnia and overall survival, Kaplan-Meier (KM) curves were plotted using the survminer package, based on the median overall survival. The grouping of the KM curves was based on comparing the insomnia scores of the cancer samples, with patients categorized into high and low insomnia score groups. A P-value < 0.05 for the KM curve was considered to indicate that the insomnia score had prognostic value for the respective cancer.

### 2.7 Pan-cancer analysis in a GDC Pan-Cancer cohort

Samples common to the basic phenotype, survival data, and gene expression data of the GDC Pan-Cancer cohort were identified. The gene expression profiles of these samples were obtained by merging the downloaded gene expression data of the GDC Pan-Cancer cohort. Subsequent analyses followed the same methodology as the pan-cancer analysis of the TCGA Pan-Cancer cohort.

## 3. Results

### 3.1 Acquisition of Insomnia Differential Genes and Related Genes

To explore the association between insomnia and cancer, we initially identified the intersection of gene symbols shared among the three datasets: GSE208668, GSE40562, and TCGA Pan-Cancer (Figure 1a). A total of 15,703 gene symbols were found common across these datasets.

**Fig 1.**
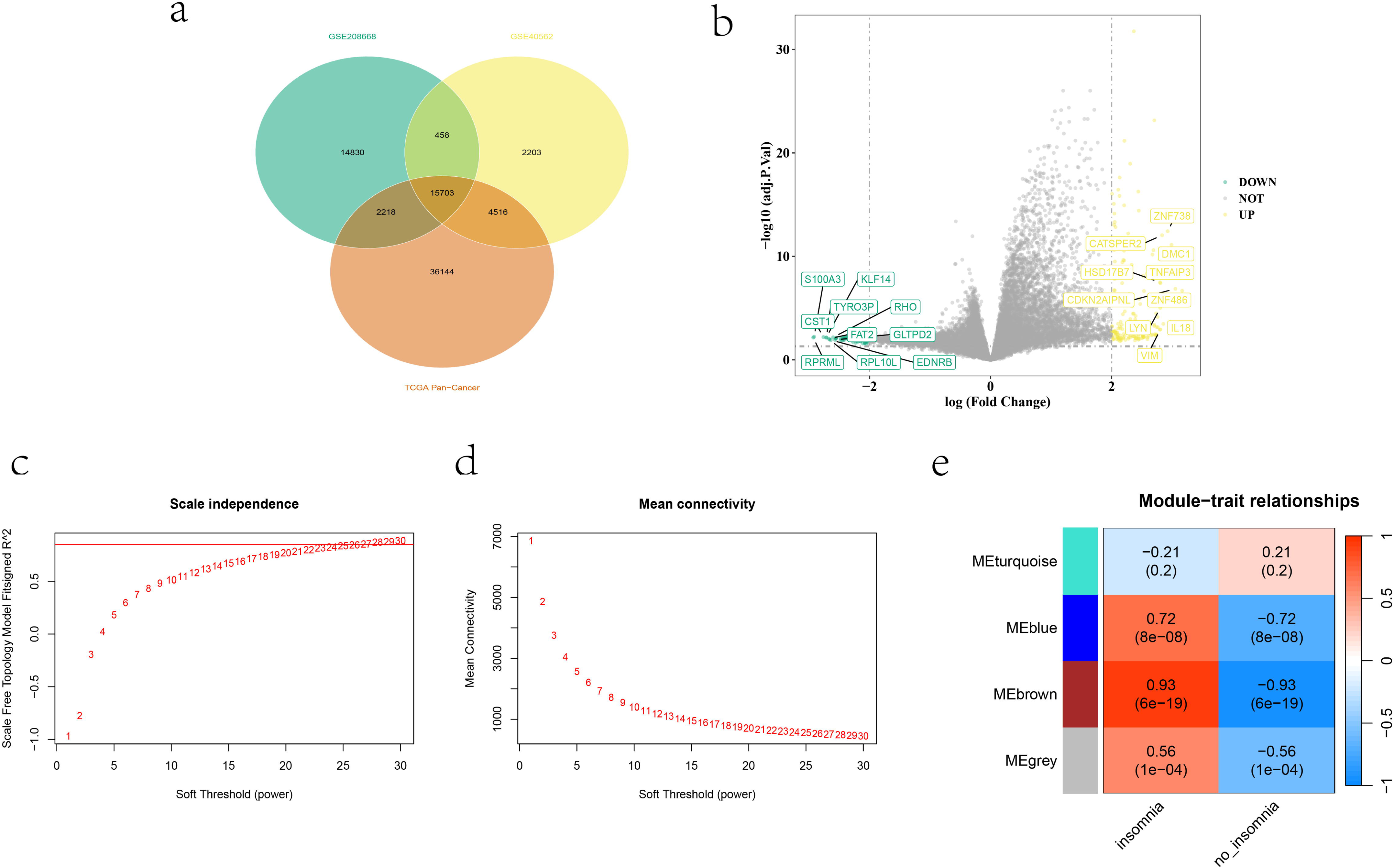
Acquisition of Insomnia Differential Genes and Related Genes. a, Gene symbols shared by GSE208668, GSE40562 and TCGA Pan-Cancer. b, Differential analysis (log_2_FoldChange>2 and FDR<0.05 marking significantly different genes) for the GSE208668 dataset, with plots demonstrating the 10 up-regulated (labeled yellow) genes and 10 down-regulated (labeled green) genes with the most significant differences. c, Scale independence for WGCNA. d, Mean connectivity for WGCNA. e, Module-trait relationships for WGCNA.

Subsequently, focusing on GSE208668, we conducted differential expression analysis between insomnia and non-insomnia conditions, visualized in the volcano plot (Figure 1b), which identified 260 genes with significant differences (Supplementary file2).

For further investigation of insomnia-related genes, we performed WGCNA using GSE208668 samples as the phenotype. Setting a soft threshold of 27 and a non-scale R2 of 0.85, we identified four distinct modules (Figure 1c-d). Notably, the MEblue and MEbrown modules showed the highest correlation coefficients and significance, leading us to select genes from these modules as insomnia-related (Figure 1e).

### 3.2 Construction and validation of an insomnia score

Using WGCNA, we identified 3,632 insomnia-related genes from GSE208668 samples (Figure 2a). Further refining this set, we identified 95 insomnia candidate genes by intersecting these 3,632 genes with those showing significant differential expression related to insomnia (Figure 2a). Subsequently, KEGG and GO enrichment analyses of these insomnia candidate genes revealed the top five significantly enriched GO terms, including leukocyte proliferation, leukocyte cell-cell adhesion, negative regulation of T cell mediated cytotoxicity, regulation of immune effector process, and immune response-inhibiting signal transduction (Figure 2b). Pathway enrichment analysis highlighted significant involvement in the B cell receptor signaling pathway and osteoclast differentiation (Figure 2c), suggesting a predominant role in immune-related functions and processes. From the insomnia candidate genes, we selected three key genes using LASSO: PTMA, NLRP8, and CCBE1 (Figure 2d-e). The coefficients obtained from LASSO analysis were as follows: Coefficient_PTMA_ = - 0.8039667, Coefficient_NLRP8_ = -3.1975230, and Coefficient_CCBE1_ = - 0.4957560 (Table 1). These coefficients were integrated into the riskScore formula (riskScore = -0.8039667 * Expression_PTMA_ + -3.1975230 * Expression_NLRP8_ + -0.4957560 * Expression_CCBE1_) to compute the insomnia score for each sample, enabling prediction and assessment of insomnia risk (Figure 2f-g).

**Fig 2.**
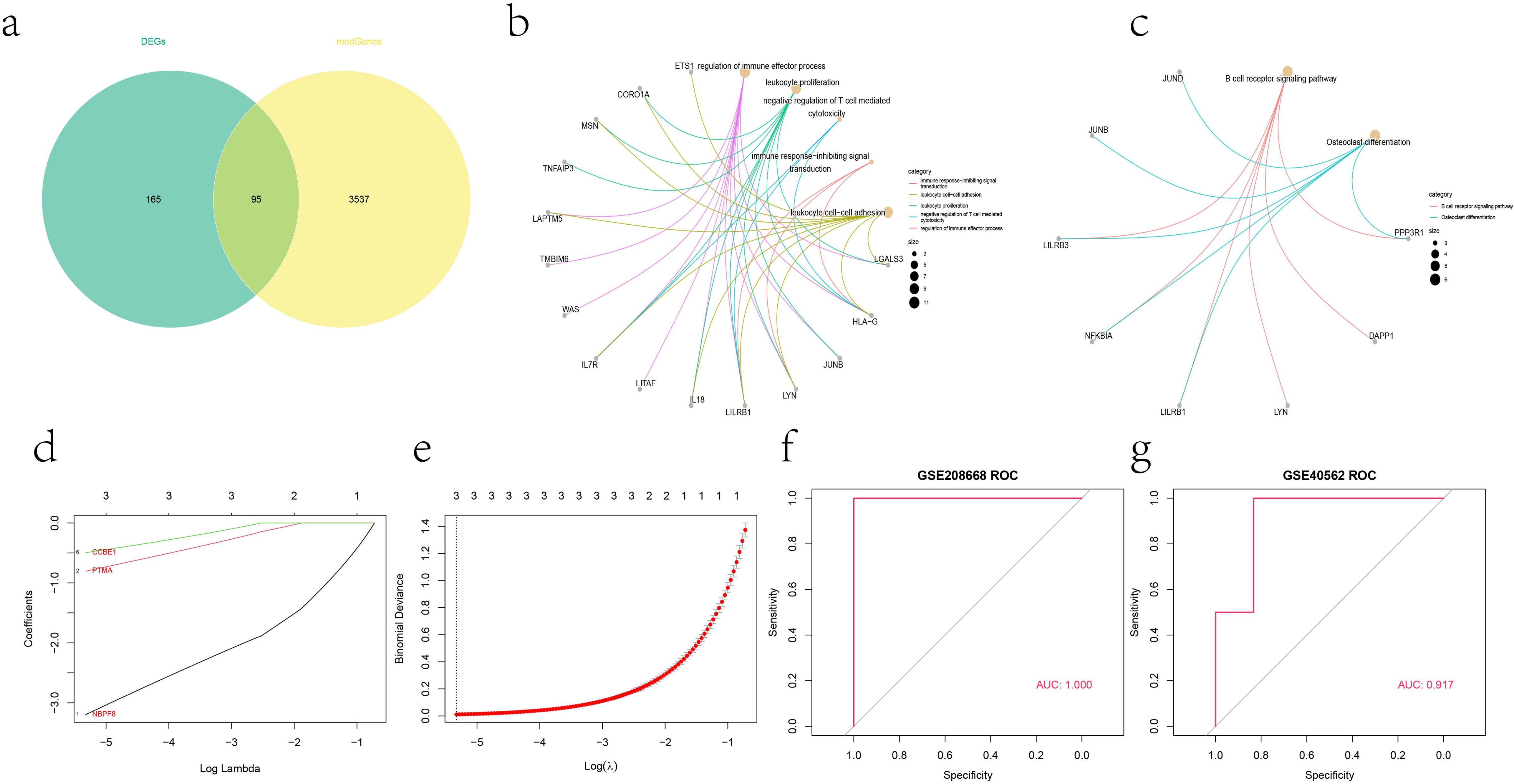
Construction and validation of an insomnia risk score. a, Differential versus modular gene veen plots, DEGs: Differently expressed genes, modGenes: MEblue module gene. b, GO enrichment analysis of insomnia candidate genes, cnetplot showing the most significantly enriched GO terms and the insomnia candidate genes they contain. c, KEGG enrichment analysis of insomnia candidate genes, cnetplot showing the most significantly enriched pathways and the insomnia candidate genes they contain. d, Coefficient path plot. e, Cross-validation error curve. f, ROC analysis of the GSE208668 dataset. g, ROC analysis of the GSE40562 dataset.

**Table 1.** Insomnia key genes and their coefficient correspondence.

| gene | Coefficient |
| --- | --- |
| PTMA | -0.8039667 |
| NLRP8 | -3.1975230 |
| CCBE1 | -0.4957560 |

In predicting insomnia traits for GSE208668 and GSE40562 datasets, the area under the curve (AUC) values were 1 and 0.917, respectively (Figure 2f-g), demonstrating excellent predictive performance of the insomnia score. Moreover, PTMA, NLRP8, and CCBE1 individually exhibited strong predictive capabilities for insomnia in both datasets (Figure 3).

**Fig 3.**
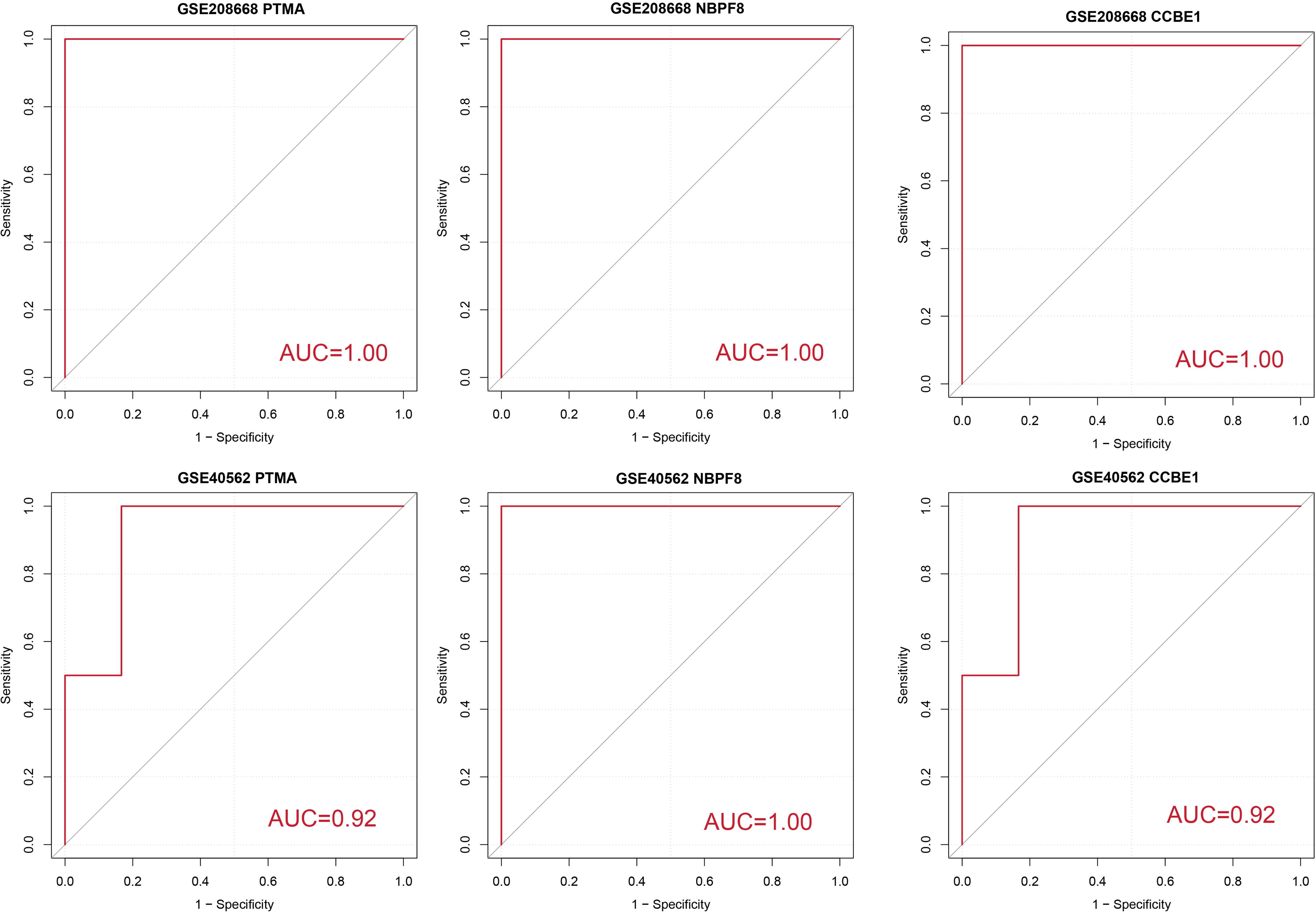
AUC values for ROC analysis of PTMA, NLRP8 and CCBE1 in the GSE208668 and GSE40562 datasets.

### 3.3 TCGA Pan-Cancer cohort pan-cancer analysis shows diagnostic value of insomnia score in multiple cancers

Insomnia has been reported to be used to predict cancer occurrence[24–29], but the mechanisms involved are currently unclear. To investigate the association between insomnia and cancer occurrence, we utilized pan-cancer data from the TCGA Pan-Cancer cohort obtained from the UCSC Xena database. We calculated the insomnia score for each sample in this dataset and visualized the differences in insomnia scores between cancerous and normal tissues across 33 types of cancers using box-and-whisker plots (Figure 4a). Subsequently, we evaluated the diagnostic performance of the insomnia score for these cancers using ROC analysis (Figure 4b).

**Fig 4.**
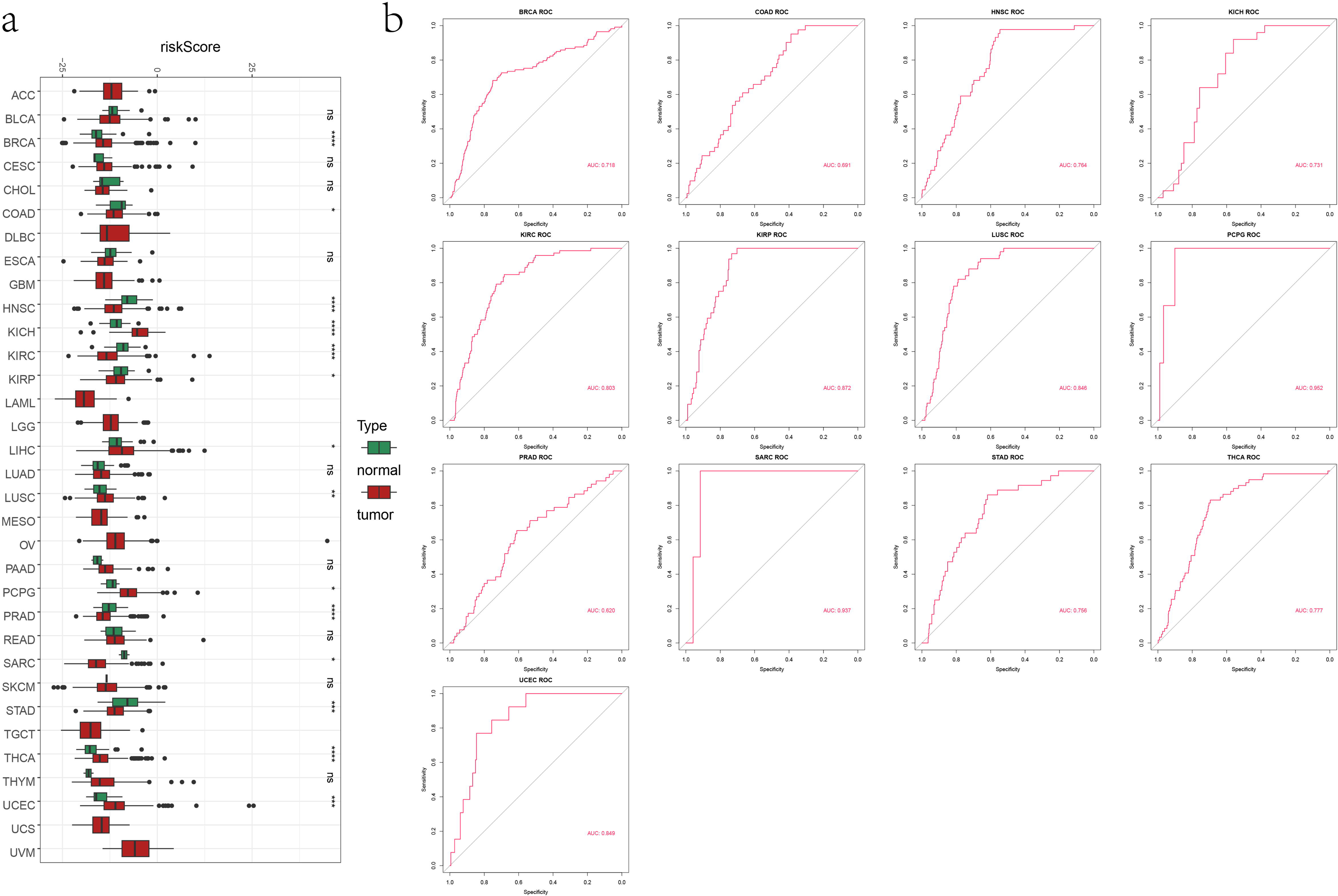
Insomnia score have diagnostic value in a variety of cancers. A, Wilcoxon rank-sum tests for tumor samples and normal samples from different cancers in the TCGA Pan-Cancer cohort, riskScore: insomnia score.Green indicates normal tissue and red indicates tumor tissue, *P < 0.05; **P < 0.01; ***P < 0.001, ****P < 0.0001,ns represents no significance. B, ROC analysis of 33 cancers from the TCGA Pan-Cancer cohort.

Our analysis revealed significant differences in insomnia scores between paired cancerous and normal tissues in several TCGA cancers (BRCA, COAD, HNSC, KICH, KIRC, KIRP, LUSC, PCPG, PRAD, SARC, STAD, THCA, and UCEC; P-value < 0.05) (Figure 4a). Furthermore, we identified cancers where the insomnia score demonstrated excellent diagnostic performance (AUC > 0.7), including BRCA, HNSC, KICH, KIRC, KIRP, LUSC, PCPG, SARC, STAD, THCA, and UCEC (Figure 4b).

### 3.4 Insomnia score as a prognostic factor for multiple cancers in the TCGA Pan-Cancer cohort

Insomnia presents as a complication in many cancer patients, prompting investigation into whether improvements in insomnia can predict better outcomes for these patients. In this study, we conducted univariate Cox proportional hazards regression analysis of overall survival and plotted Kaplan-Meier (KM) curves for 33 cancers in the TCGA Pan-Cancer cohort (Figure 5) to assess whether insomnia can serve as a prognostic factor for cancer.

**Fig 5.**
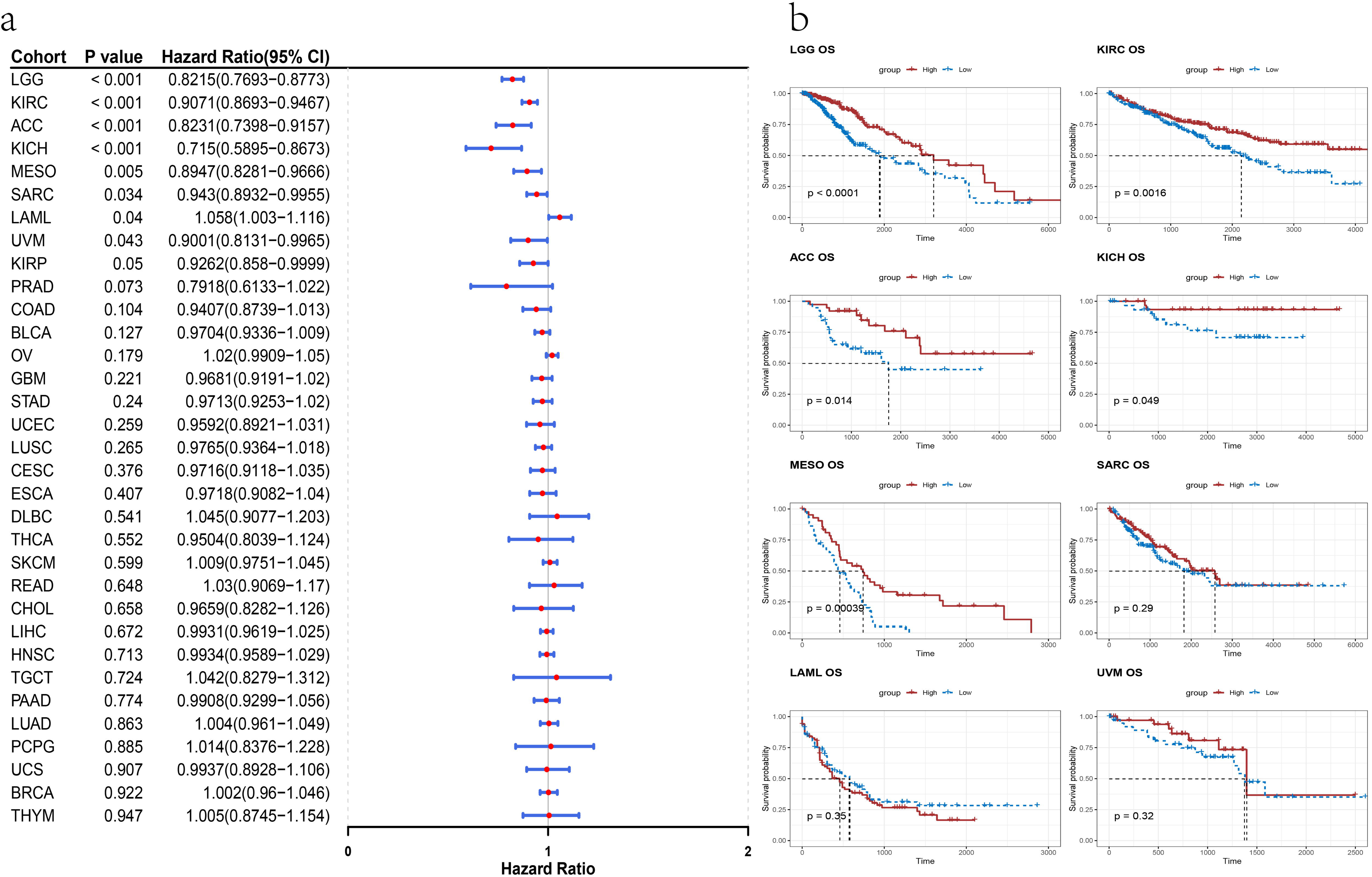
Insomnia score can be a prognostic factor for many cancers. A, Univariate Cox proportional hazards regression analysis of insomnia score in the TCGA Pan-Cancer cohort. B, Overall survival KM curve analysis of cancers in the TCGA Pan-Cancer cohort in which overall survival was significantly affected by insomnia score. Red indicates survival curves with insomnia score above the mean and blue indicates survival curves with insomnia score below the mean. Taking the median overall survival to plot the KM curve.

In the results of the univariate Cox proportional hazards regression analysis of the TCGA Pan-Cancer cohort, insomnia score significantly influenced overall survival in UVM, LGG, ACC, MESO, KIRC, KICH, SARC, and LAML (P-value < 0.05). Notably, overall survival was positively correlated with insomnia score only for LAML, while it was negatively correlated for the other cancers (Figure 5a). To validate these findings, we selected cancers where overall survival was significantly affected by insomnia score for KM analysis, demonstrating differences in overall survival times between groups with high and low insomnia scores, particularly in LGG, ACC, MESO, KIRC, and KICH, where higher insomnia scores were associated with shorter overall survival times (P-value < 0.05) (Figure 5b). These results indicate that insomnia score serves as a prognostic factor for LGG, ACC, MESO, KIRC, and KICH conditions. Specifically, insomnia score exhibits both diagnostic and prognostic value in KIRC, where a lower score may suggest higher prevalence and poorer prognosis (Figures 4 and 5). However, such a pattern was not observed for the KICH cohort, indicating a different relationship between insomnia score and diagnostic survival compared to KIRC (Figures 4 and 5).

### 3.5 Analysis of the diagnostic and prognostic value of the insomnia score in a GDC Pan-Cancer cohort

To ensure robustness, we validated and extended our analysis using the GDC Pan-Cancer dataset in parallel with the TCGA Pan-Cancer data (Figures 6 and 7). After computing the insomnia score for each sample in the GDC Pan-Cancer cohort, significant intergroup differences in insomnia scores were observed between cancer and control groups in TCGA-BRCA, TCGA-CESC, TCGA-CHOL, TCGA-COAD, TCGA-KICH, TCGA-KIRC, TCGA-KIRP, TCGA-LIHC, TCGA-LUAD, TCGA-PCPG, TCGA-PRAD, TCGA-STAD, TCGA-THCA, and TCGA-THYM (P-value < 0.05) (Figure 6a). ROC analysis further demonstrated the diagnostic capability of insomnia score in predicting the onset of TCGA-BRCA, TCGA-CESC, TCGA-CHOL, TCGA-KICH, TCGA-LIHC, TCGA-PCPG, TCGA-STAD, and TCGA-THYM with high performance (AUC > 0.7) (Figure 6b). Hence, insomnia score exhibits diagnostic value across these cancers in the GDC Pan-Cancer cohort.

**Fig 6.**
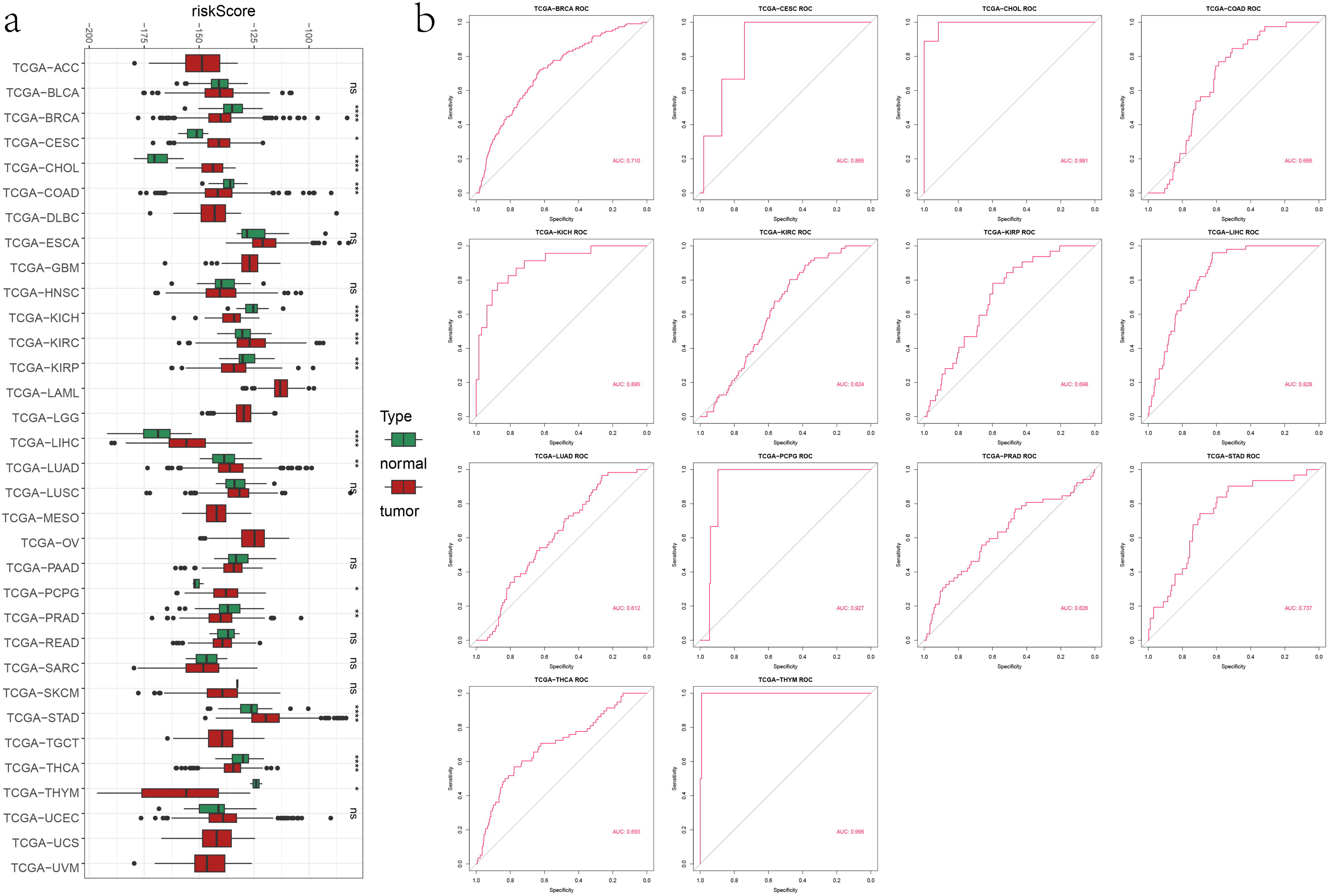
Analysis of the diagnostic and prognostic value of the insomnia score in a GDC Pan-Cancer cohort. A, Wilcoxon rank-sum tests for tumor samples and normal samples from different cancers in the TCGA Pan-Cancer cohort, riskScore: insomnia score.Green indicates normal tissue and red indicates tumor tissue, *P < 0.05; **P < 0.01; ***P < 0.001, ****P < 0.0001,ns represents no significance. B, ROC analysis of cancers in the GDC Pan-Cancer cohort in which insomnia score were significantly different between cancer and control groups.

**Fig 7.**
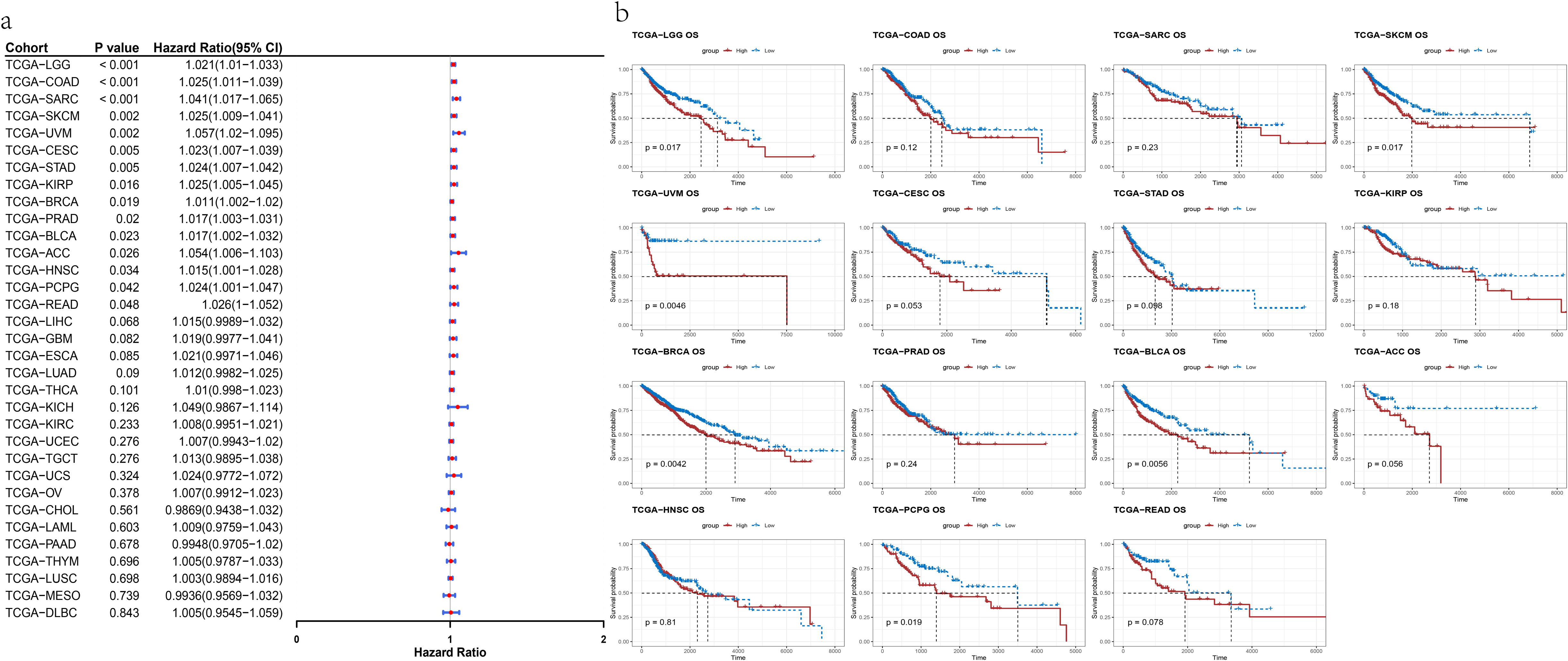
Analysis of the diagnostic and prognostic value of the insomnia score in a GDC Pan-Cancer cohort. A, Univariate Cox proportional hazards regression analysis of the GDC Pan-Cancer cohort. B, KM curves for cancers in the GDC Pan-Cancer cohort in which overall survival was significantly affected by insomnia score. Red indicates survival curves with insomnia score above the mean and blue indicates survival curves with insomnia score below the mean. Taking the median overall survival to plot the KM curve.

Subsequently, we assessed the prognostic value of insomnia score in cancer samples from the GDC Pan-Cancer cohort (Figure 7). Univariate Cox proportional hazards regression analyses revealed that insomnia score significantly influenced overall survival in TCGA-LGG, TCGA-COAD, TCGA-BRCA, TCGA-SARC, TCGA-SKCM, TCGA-UVM, TCGA-CESC, TCGA-STAD, TCGA-KIRP, TCGA-PRAD, TCGA-ACC, TCGA-HNSC, TCGA-BLCA, TCGA-READ, and TCGA-PCPG (P < 0.05), with higher insomnia scores correlating with poorer overall survival rates across these cancers (Figure 7a). Kaplan-Meier curves further illustrated significant differences in overall survival times between high and low insomnia score groups in TCGA-LGG, TCGA-BRCA, TCGA-UVM, TCGA-SKCM,

TCGA-BLCA, and TCGA-PCPG, where lower insomnia scores were associated with longer overall survival (P-value < 0.05) (Figure 7b). These findings indicate that insomnia score serves as a prognostic indicator in LGG, BRCA, UVM, SKCM, BLCA, and PCPG populations.

Moreover, insomnia score exhibits both diagnostic and prognostic value specifically for BRCA and PCPG in the GDC Pan-Cancer cohort, where higher scores are associated with increased likelihood of cancer prevalence and poorer prognosis.

## 4. Discussion

In this study, we identified insomnia key genes from the GSE208668 insomnia cohort and utilized machine learning techniques to construct an insomnia score. Upon validation and evaluation, both the insomnia core genes and insomnia score demonstrated exceptional diagnostic ability for insomnia symptoms in the GSE208668 and GSE40562 datasets (Figure 3). Notably, according to UniProt information (https://www.uniprot.org/)[44], the roles of NBPF8, PTMA and CCBE1 in insomnia have been less characterized, warranting further in-depth investigation into their mechanisms in insomnia.

Subsequently, we analyzed the diagnostic and prognostic functions of insomnia score in two pan-cancer cohorts, TCGA Pan-Cancer and GDC Pan-Cancer, to assess its utility in cancer conditions. Our findings revealed that insomnia score possesses diagnostic and prognostic values across numerous cancers, indicating a significant correlation between insomnia and cancer (Figures 4-7).

This study breaks new ground by exploring the association between insomnia and cancer using gene expression data, complementing existing research on insomnia-cancer associations at the gene expression level. Insomnia scores demonstrated diagnostic and prognostic value across pan-cancer cohorts, underscoring their potential clinical relevance. However, further experimental validation of these findings is essential.

Despite these advancements, several limitations should be noted. The primary shortcomings stem from data gaps, including the limited availability of comprehensive insomnia datasets and pan-cancer data covering diverse populations. Most existing insomnia datasets focus on adults, with scant data available for adolescents, despite the significant issue of insomnia in this age group [5,45]. Our analysis of the TARGET Pan-Cancer cohort, which includes minors, did not reveal significant associations between insomnia score and cancers (Supplementary file3). This may be attributed to the relatively sparse nature of the minor pan- cancer cohort compared to the TCGA Pan-Cancer and GDC Pan-Cancer datasets.

Insomnia is recognized as a common complication of cancer [15–20], and some studies suggest that insomnia may increase cancer risk[24–29]. The lack of recorded insomnia data in clinical information from cohorts used primarily for pan-cancer studies currently hinders the ability to determine whether insomnia is a cause or consequence of cancer at the gene expression level. Objective measurements of insomnia are crucial for achieving more accurate results[14] and advancing personalized and targeted care.

In conclusion, our findings highlight not only the better predictability of insomnia scores for insomnia, but also their potential in the diagnosis and prognosis of various cancers, revealing associations between insomnia and cancer at the gene expression level. Although our data suggest a promising application of the insomnia score in cancer diagnosis and prognosis, we advocate combining the insomnia score with other clinical and biomarkers to improve accuracy and reliability. Further studies are needed to validate the validity and intrinsic mechanisms of the insomnia score as an independent diagnostic and prognostic model.

## 5. Conclusion

We developed an insomnia score and confirmed its exceptional predictive capability for insomnia. Pan-cancer analyses demonstrated that this score holds diagnostic and prognostic significance across multiple cancer types, offering novel insights into the relationship between insomnia and cancer.

## Abbreviations

WGCNA: Weighted correlation network analysis
GO: Gene ontology
KEGG: Kyoto encyclopedia of genes and genomes
LASSO: Least Absolute Shrinkage and Selection Operator
PTMA: Prothymosin alpha
NLRP8: NACHT, LRR and PYD domains-containing protein 8
CCBE1: Collagen and calcium-binding EGF domain-containing protein 1
PBMCs: peripheral blood mononuclear cells
TPM: Transcripts Per Million
ROC: Receiver operating characteristic curve
AUC: Area Under Curve
KM curve: Kaplan-Meier curve
BRCA: Breast Cancer
UCEC: Uterine Corpus Endometrial Carcinoma
KIRC: Kidney Renal Clear Cell Carcinoma
HNSC: Head and Neck Squamous Cell Carcinoma
LUAD: Lung Adenocarcinoma
LGG: Low Grade Glioma
THCA: Thyroid Cancer
LUSC: Lung Squamous Cell Carcinoma
PRAD: Prostate Adenocarcinoma
SKCM: Cutaneous Melanoma
COAD: Colorectal Adenocarcinoma
OV: Ovarian Cancer
STAD: Stomach Adenocarcinoma
BLCA: Bladder Urothelial Carcinoma
LIHC: LIver Hepatocellular Carcinoma
CESC: Cervical Squamous Cell Carcinoma and Endocervical Adenocarcinoma
KIRP: Kidney Renal Papillary Cell Carcinoma
SARC: Sarcoma
ESCA: Esophageal Carcinoma
PCPG: Pheochromocytoma and Paraganglioma
PAAD: Pancreatic Adenocarcinoma
GBM: Glioblastoma Multiforme
READ: Rectal Adenocarcinoma
LAML: Acute Myeloid Leukemia
TGCT: Testicular Germ Cell Tumor
THYM: Thymoma
MESO: Malignant Mesothelioma
UVM: Uveal Melanoma
ACC: Adrenocortical carcinoma
KICH: Kidney Chromophobe
USC: Uterine Carcinosarcoma
DLBC: Diffuse Large B-cell Lymphoma
CHOL: Cholangiocarcinoma
TPM: Transcripts per million
FDR: False Discovery Rate
OS: Overall Survival.

## Data availability

This paper analyzes existing, publicly available data. Microarray are openly available in GEO (https://www.ncbi.nlm.nih.gov/geo/),accession numbers [GSE208668 and GSE40562]. Gene expression, clinical information and survival information data for 33 cancers from TCGA in UCSC Xena(https://xenabrowser.net/datapages/).Any additional information needed to reanalyze the data reported in this paper can be obtained from the primary contact upon request.

## Acknowledgments

We thank the Gene Expression Omnibus (GEO) and The Cancer Genome Atlas (TCGA) Database for sharing a large amount of data. Thanks to UCSC Xena for collecting and organizing the pan-cancer data.

## Appendix. Supplementary materials

Supplementary file1

Overlap of GDC Pan-Cancer and TARGET Pan-Cancer samples in the UCSC Xena database.

Supplementary file2

GSE208668 differential analysis results

Supplementary file3

Pan-cancer analysis of the TARGET Pan-Cancer Cohort at UCSC Xena.

## References

[1] E.L. Sutton, Insomnia, Ann Intern Med. 174 (2021) ITC33-ITC48. 10.7326/AITC202103160.

[2] D. Riemann, C. Nissen, L. Palagini, A. Otte, M.L. Perlis, K. Spiegelhalder, The neurobiology, investigation, and treatment of chronic insomnia, Lancet Neurol. 14 (2015) 547–558. 10.1016/S1474-4422(15)00021-6.

[3] D.J. Buysse, Insomnia, JAMA. 309 (2013) 706-716. 10.1001/jama.2013.193.

[4] D. Patel, J. Steinberg, P. Patel, Insomnia in the Elderly: A Review, J Clin Sleep Med. 14 (2018) 1017–1024. 10.5664/jcsm.7172.

[5] M. Himelfarb, J.P. Shatkin, Pediatric Insomnia, Child Adolesc Psychiatr Clin N Am. 30 (2021) 117–129. 10.1016/j.chc.2020.08.004.

[6] Z.L. Cohen, P.M. Eigenberger, K.M. Sharkey, M.L. Conroy, K.M. Wilkins, Insomnia and Other Sleep Disorders in Older Adults, Psychiatr Clin North Am. 45 (2022) 717–734. 10.1016/j.psc.2022.07.002.

[7] M. Jansson-Frojmark, K. Lindblom, A bidirectional relationship between anxiety and depression, and insomnia? A prospective study in the general population, J Psychosom Res. 64 (2008) 443–449. 10.1016/j.jpsychores.2007.10.016.

[8] W.R. Pigeon, M. Hegel, J. Unutzer, M.Y. Fan, M.J. Sateia, J.M. Lyness, C. Phillips, M.L. Perlis, Is insomnia a perpetuating factor for late-life depression in the IMPACT cohort?, Sleep. 31 (2008) 481–488. 10.1093/sleep/31.4.481.

[9] L. Palagini, R.M. Bruno, A. Gemignani, C. Baglioni, L. Ghiadoni, D. Riemann, Sleep loss and hypertension: a systematic review, Curr Pharm Des. 19 (2013) 2409–2419. 10.2174/1381612811319130009.

[10] A.N. Vgontzas, D. Liao, E.O. Bixler, G.P. Chrousos, A. Vela-Bueno, Insomnia with objective short sleep duration is associated with a high risk for hypertension, Sleep. 32 (2009) 491–497. 10.1093/sleep/32.4.491.

[11] L.E. Laugsand, L.J. Vatten, C. Platou, I. Janszky, Insomnia and the risk of acute myocardial infarction: a population study, Circulation. 124 (2011) 2073–2081. 10.1161/CIRCULATIONAHA.111.025858.

[12] W.M. Troxel, D.J. Buysse, K.A. Matthews, K.E. Kip, P.J. Strollo, M. Hall, O. Drumheller, S.E. Reis, Sleep symptoms predict the development of the metabolic syndrome, Sleep. 33 (2010) 1633–1640. 10.1093/sleep/33.12.1633.

[13] D.J. Gottlieb, N.M. Punjabi, A.B. Newman, H.E. Resnick, S. Redline, C.M. Baldwin, F.J. Nieto, Association of sleep time with diabetes mellitus and impaired glucose tolerance, Arch Intern Med. 165 (2005) 863–867. 10.1001/archinte.165.8.863.

[14] J. Fernandez-Mendoza, A.N. Vgontzas, Insomnia and its impact on physical and mental health, Curr Psychiatry Rep. 15 (2013) 418. 10.1007/s11920-013-0418-8.

[15] R.R. Induru, D. Walsh, Cancer-related insomnia, Am J Hosp Palliat Care. 31 (2014) 777–785. 10.1177/1049909113508302.

[16] A. Kwak, J. Jacobs, D. Haggett, R. Jimenez, J. Peppercorn, Evaluation and management of insomnia in women with breast cancer, Breast Cancer Res Treat. 181 (2020) 269–277. 10.1007/s10549-020-05635-0.

[17] O.G. Palesh, J.A. Roscoe, K.M. Mustian, T. Roth, J. Savard, S. Ancoli-Israel, C. Heckler, J.Q. Purnell, M.C. Janelsins, G.R. Morrow, Prevalence, demographics, and psychological associations of sleep disruption in patients with cancer: University of Rochester Cancer Center-Community Clinical Oncology Program, J Clin Oncol. 28 (2010) 292-298. 10.1200/JCO.2009.22.5011.

[18] K. Schieber, A. Niecke, F. Geiser, Y. Erim, C. Bergelt, A. Buttner-Teleaga, I. Maatouk, B. Stein, M. Teufel, M. Wickert, A. Wuensch, J. Weis, The course of cancer-related insomnia: don’t expect it to disappear after cancer treatment, Sleep Med. 58 (2019) 107–113. 10.1016/j.sleep.2019.02.018.

[19] A.K. Wong, D.R.Y. Wang, D. Marco, B. Le, J. Philip, Prevalence, Severity, and Predictors of Insomnia in Advanced Colorectal Cancer, Journal of Pain and Symptom Management. 66 (2023) e335–e342. 10.1016/j.jpainsymman.2023.05.020.

[20] J. Zhang, Z. Zhang, S. Huang, X. Qiu, L. Lao, Y. Huang, Z.J. Zhang, Acupuncture for cancer-related insomnia: A systematic review and meta-analysis, Phytomedicine. 102 (2022) 154160. 10.1016/j.phymed.2022.154160.

[21] L. Fleming, S. Gillespie, C.A. Espie, The development and impact of insomnia on cancer survivors: a qualitative analysis, Psychooncology. 19 (2010) 991–996. 10.1002/pon.1652.

[22] J. Savard, H. Ivers, M.H. Savard, C.M. Morin, Cancer treatments and their side effects are associated with aggravation of insomnia: Results of a longitudinal study, Cancer. 121 (2015) 1703–1711. 10.1002/cncr.29244.

[23] J. Zhang, Z. Qin, T.H. So, T.Y. Chang, S. Yang, H. Chen, W.F. Yeung, K.F. Chung, P.Y. Chan, Y. Huang, S. Xu, C.Y. Chiang, L. Lao, Z.J. Zhang, Acupuncture for chemotherapy-associated insomnia in breast cancer patients: an assessor-participant blinded, randomized, sham-controlled trial, Breast Cancer Res. 25 (2023) 49. 10.1186/s13058-023-01645-0.

[24] Z. Huo, F. Ge, C. Li, H. Cheng, Y. Lu, R. Wang, Y. Wen, K. Yue, Z. Pan, H. Peng, X. Wu, H. Liang, J. He, W. Liang, Genetically predicted insomnia and lung cancer risk: a Mendelian randomization study, Sleep Med. 87 (2021) 183–190. 10.1016/j.sleep.2021.06.044.

[25] L. Palagini, M. Miniati, L. Massa, F. Folesani, D. Marazziti, L. Grassi, D. Riemann, Insomnia and circadian sleep disorders in ovarian cancer: Evaluation and management of underestimated modifiable factors potentially contributing to morbidity, J Sleep Res. 31 (2022) e13510. 10.1111/jsr.13510.

[26] A. Sen, S. Opdahl, L.B. Strand, L.J. Vatten, L.E. Laugsand, I. Janszky, Insomnia and the Risk of Breast Cancer: The HUNT Study, Psychosom Med. 79 (2017) 461–468. 10.1097/PSY.0000000000000417.

[27] T. Shi, M. Min, C. Sun, Y. Zhang, M. Liang, Y. Sun, Does insomnia predict a high risk of cancer? A systematic review and meta-analysis of cohort studies, J Sleep Res. 29 (2020) e12876. 10.1111/jsr.12876.

[28] H. Wang, B.M. Reid, R.C. Richmond, J.M. Lane, R. Saxena, B.D. Gonzalez, B.L. Fridley, S. Redline, S.S. Tworoger, X. Wang, Impact of insomnia on ovarian cancer risk and survival: a Mendelian randomization study, EBioMedicine. 104 (2024) 105175. 10.1016/j.ebiom.2024.105175.

[29] K. Yoon, C.M. Shin, K. Han, J.H. Jung, E.H. Jin, J.H. Lim, S.J. Kang, Y.J. Choi, D.H. Lee, Risk of cancer in patients with insomnia: Nationwide retrospective cohort study (2009-2018), PLoS One. 18 (2023) e0284494. 10.1371/journal.pone.0284494.

[30] P. Langfelder, S. Horvath, WGCNA: an R package for weighted correlation network analysis, BMC Bioinformatics. 9 (2008) 559. 10.1186/1471-2105-9-559.

[31] J. Friedman, T. Hastie, R. Tibshirani, Regularization Paths for Generalized Linear Models via Coordinate Descent, J Stat Softw. 33 (2010) 1–22.

[32] J. Chen, M. Sun, C. Chen, B. Jiang, Y. Fang, Identification of hub genes and their correlation with infiltration of immune cells in MYCN positive neuroblastoma based on WGCNA and LASSO algorithm, Front Immunol. 13 (2022) 1016683. 10.3389/fimmu.2022.1016683.

[33] F. Jiang, H. Zhou, H. Shen, Identification of Critical Biomarkers and Immune Infiltration in Rheumatoid Arthritis Based on WGCNA and LASSO Algorithm, Front Immunol. 13 (2022) 925695. 10.3389/fimmu.2022.925695.

[34] Y. Jin, S. Huang, Z. Wang, Identify and validate RUNX2 and LAMA2 as novel prognostic signatures and correlate with immune infiltrates in bladder cancer, Front Oncol. 13 (2023) 1191398. 10.3389/fonc.2023.1191398.

[35] H. Song, Y. Ge, J. Xu, R. Shen, P.C. Zhang, G.Q. Wang, B. Liu, Identification and validation of novel signature associated with hepatocellular carcinoma prognosis using Single-cell and WGCNA analysis, Int J Med Sci. 20 (2023) 870–887. 10.7150/ijms.79274.

[36] Z. Wang, J. Liu, Y. Wang, H. Guo, F. Li, Y. Cao, L. Zhao, H. Chen, Identification of Key Biomarkers Associated with Immunogenic Cell Death and Their Regulatory Mechanisms in Severe Acute Pancreatitis Based on WGCNA and Machine Learning, Int J Mol Sci. 24 (2023). 10.3390/ijms24033033.

[37] W. Li, Y. Gao, X. Jin, H. Wang, T. Lan, M. Wei, W. Yan, G. Wang, Z. Li, Z. Zhao, X. Jiang, Comprehensive analysis of N6-methylandenosine regulators and m6A-related RNAs as prognosis factors in colorectal cancer, Mol Ther Nucleic Acids. 27 (2022) 598–610. 10.1016/j.omtn.2021.12.007.

[38] D. Piber, J.H. Cho, O. Lee, D.M. Lamkin, R. Olmstead, M.R. Irwin, Sleep disturbance and activation of cellular and transcriptional mechanisms of inflammation in older adults, Brain Behav Immun. 106 (2022) 67–75. 10.1016/j.bbi.2022.08.004.

[39] C. Tian, D. Liu, W. Xiang, H.A. Kretzschmar, Q.L. Sun, C. Gao, Y. Xu, H. Wang, X.Y. Fan, G. Meng, W. Li, X.P. Dong, Analyses of the similarity and difference of global gene expression profiles in cortex regions of three neurodegenerative diseases: sporadic Creutzfeldt-Jakob disease (sCJD), fatal familial insomnia (FFI), and Alzheimer’s disease (AD), Mol Neurobiol. 50 (2014) 473–481. 10.1007/s12035-014-8758-x.

[40] M.J. Goldman, B. Craft, M. Hastie, K. Repecka, F. McDade, A. Kamath, A. Banerjee, Y. Luo, D. Rogers, A.N. Brooks, J. Zhu, D. Haussler, Visualizing and interpreting cancer genomics data via the Xena platform, Nat Biotechnol. 38 (2020) 675–678. 10.1038/s41587-020-0546-8.

[41] S. Davis, P.S. Meltzer, GEOquery: a bridge between the Gene Expression Omnibus (GEO) and BioConductor, Bioinformatics. 23 (2007) 1846–1847. 10.1093/bioinformatics/btm254.

[42] T. Wu, E. Hu, S. Xu, M. Chen, P. Guo, Z. Dai, T. Feng, L. Zhou, W. Tang, L. Zhan, X. Fu, S. Liu, X. Bo, G. Yu, clusterProfiler 4.0: A universal enrichment tool for interpreting omics data, Innovation (Camb). 2 (2021) 100141. 10.1016/j.xinn.2021.100141.

[43] X. Robin, N. Turck, A. Hainard, N. Tiberti, F. Lisacek, J.C. Sanchez, M. Muller, pROC: an open-source package for R and S+ to analyze and compare ROC curves, BMC Bioinformatics. 12 (2011) 77. 10.1186/1471-2105-12-77.

[44] C. UniProt, UniProt: the universal protein knowledgebase in 2021, Nucleic Acids Res. 49 (2021) D480–D489. 10.1093/nar/gkaa1100.

[45] M.L. Nunes, O. Bruni, Insomnia in childhood and adolescence: clinical aspects, diagnosis, and therapeutic approach, J Pediatr (Rio J). 91 (2015) S26–35. 10.1016/j.jped.2015.08.006.

