## Supplementary file3 for "Insomnia score: predictive ability of insomnia, high-diagnostic and prognostic value for cancer"

### TARGET Pan-Cancer Analysis

#### Preface

TARGET is a pan-cancer queue for children.

#### Acknowledgments

First, we thank the TARGET (Therapeutically Applicable Research to Generate Effective Treatments) program for providing quality data, and we promise that the TARGET data we use will not be used for the sole purposes of methods We thank UCSC Xena for collecting and organizing the TARGET Pan-Cancer data.

#### Methods

Download TARGET Pan-Cancer data of type TPM from the UCSC Xena database for pan-cancer analysis. Download TARGET sample phenotype to be organized into phenotype, and download TARGET donor phenotype to be organized into survival. Determine the gene expression data of TPM types of TARGET Pan-Cancer, samples with both survival and phenotype as shared samples, and exclude the gene expression data of non-shared samples, survival information and phenotype information. Due to the small number of samples, the primary_disease_code in the phenotype was excluded as Ewing sarcoma, CNS ependymoma, CNS glioblastoma

(GBM), CNS low grade glioma (LGG), CNS medulloblastoma, CNS rhabdoid tumor, CNS other, NHL, anaplastic large cell lymphoma, NHL Burkitt lymphoma (BL), Soft tissue sarcoma non rhabdomyosarcoma. primary_disease_code in the phenotype was used to determine the type of cancer analyzed, and Rhabdomyosarcoma was labeled as RMS.The above data can be self-arranged or contact the corresponding authors to provide us with the organized phenotype and survival data. TARGET Pan-Cancer analysis was performed as in the TCGA Pan-Cancer and GDC Pan-Cancer analyses, and the relevant analysis code can be obtained by contacting the corresponding author, and the code required for the analysis will be provided upon reasonable request.

Abbreviations

AML:Acute Myeloid Leukemia; ALL　Acute Lymphoblastic Leukemia; NBL　Neuroblastoma.

Notes

TARGET Pan-Cancer data were processed as described above, as well as not including normal tissue samples, so analyses related to assessing diagnostic value were not performed and prognostic analyses were performed directly.

Results

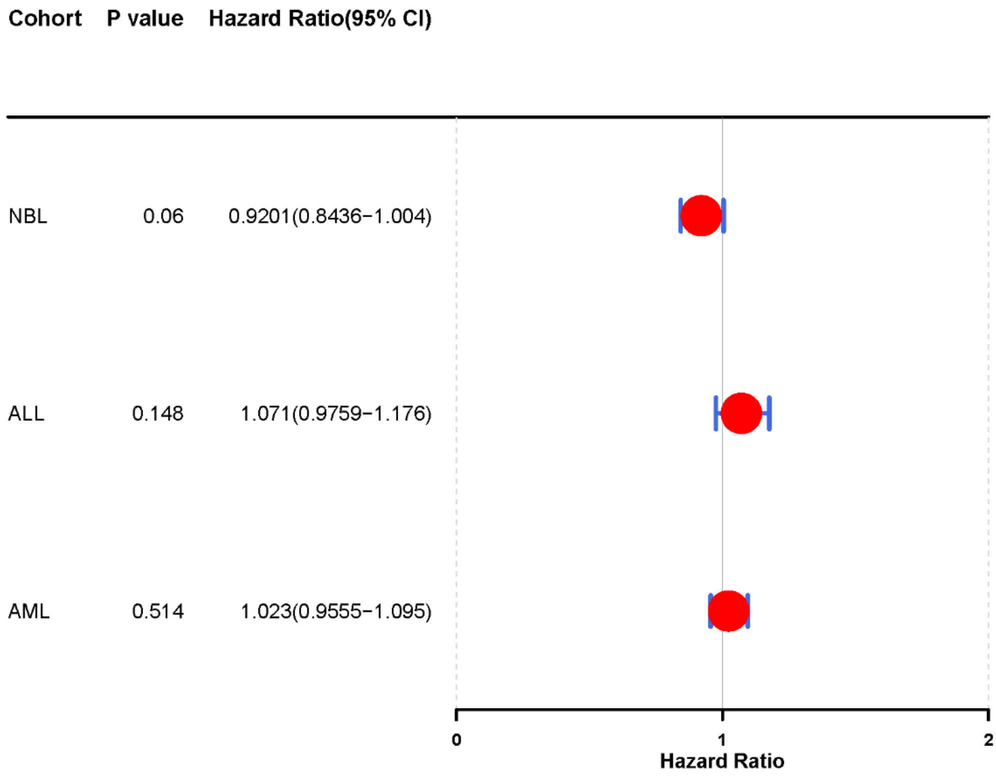


Figure1.TARGET Pan-Cancer univariate Cox proportional hazards regression analysis.

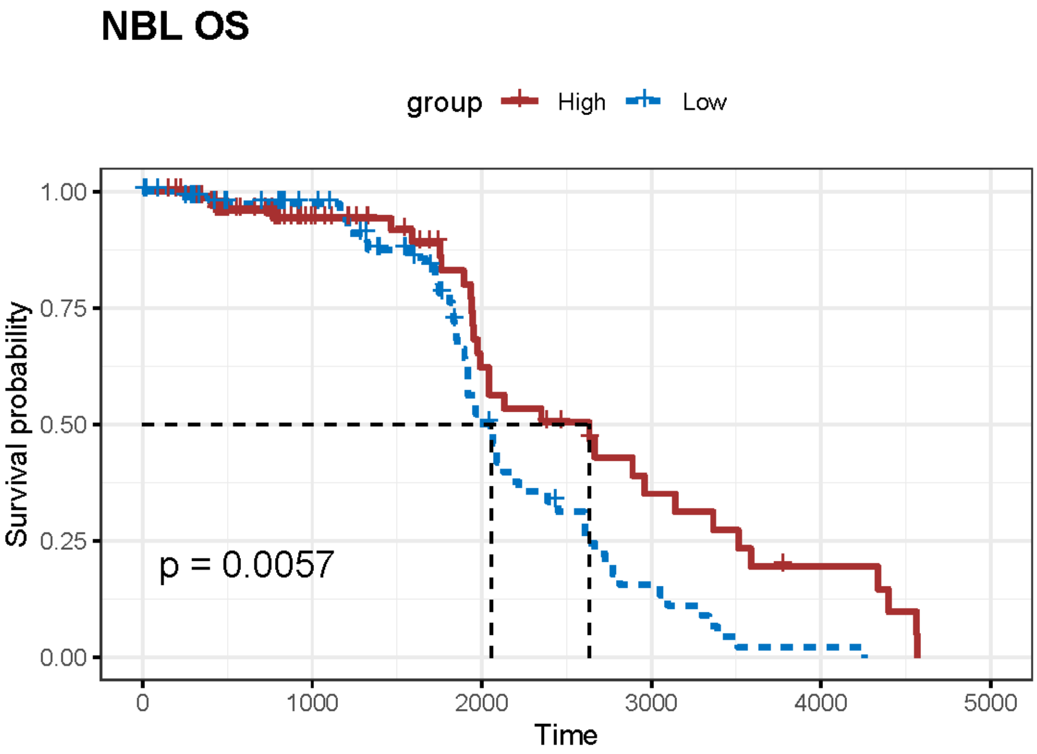

Figure2.Kaplan-Meier curve for NBL (extracted median total survival time plotted).

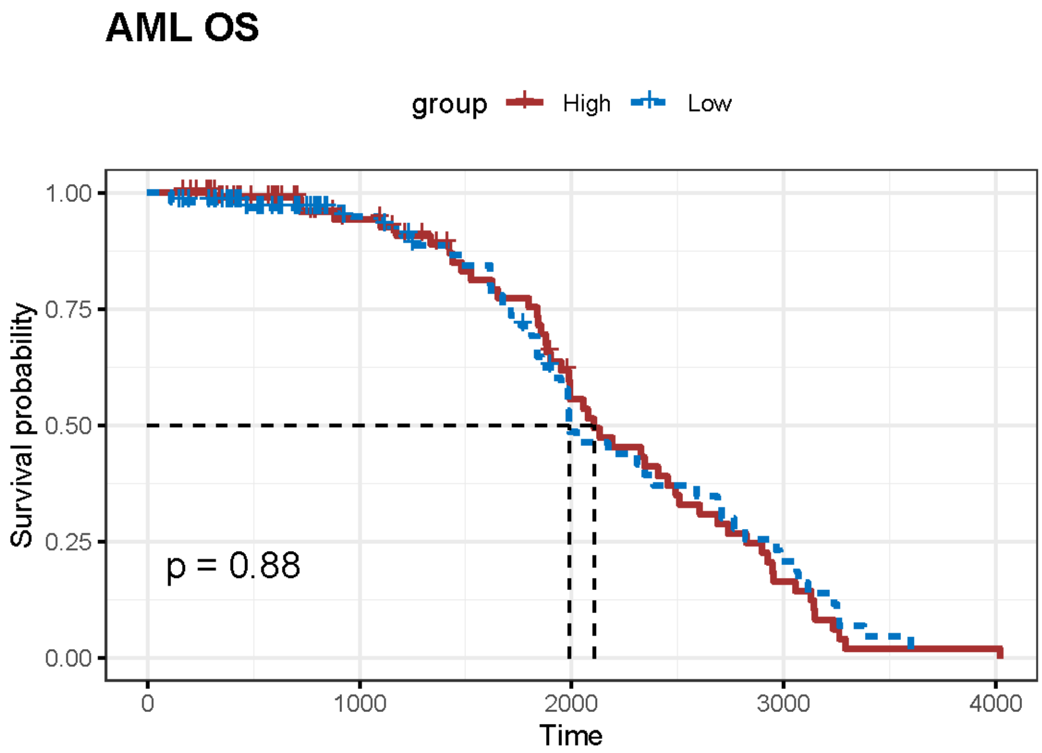


Figure3.Kaplan-Meier curve for AML (extracted median total survival time plotted).

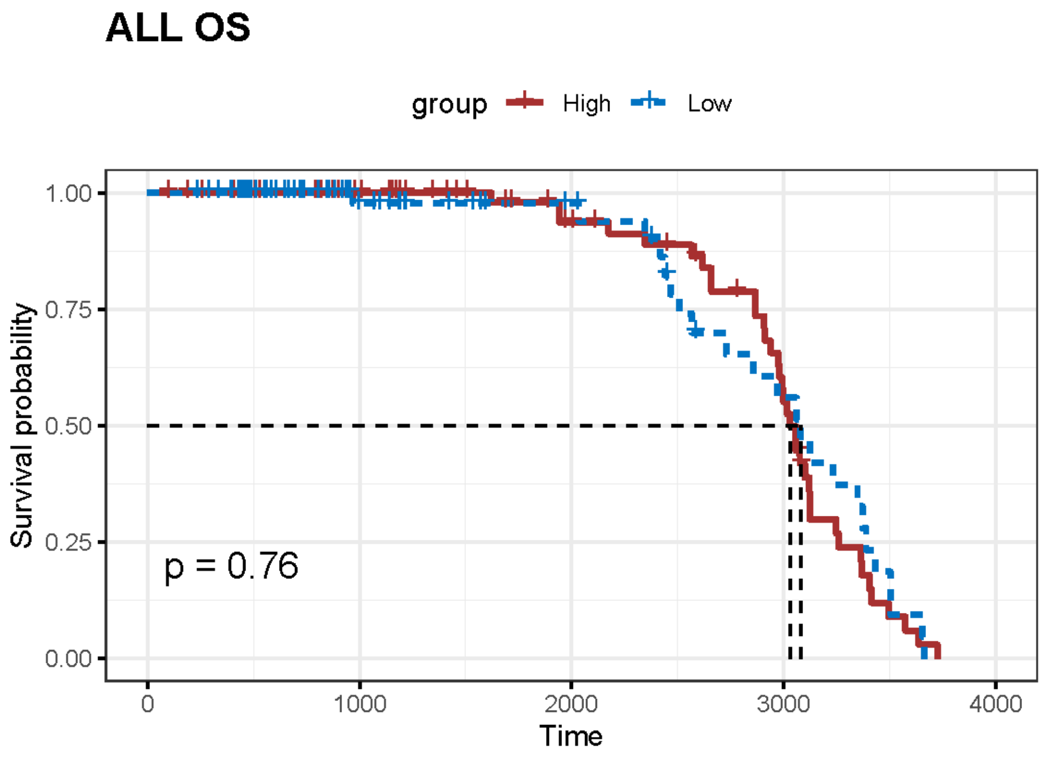


Figure 2 Kaplan-Meier curve for ALL (extracted median total survival time plotted).
